# The role of HMGB1 in invasive *Candida albicans* infection

**DOI:** 10.1101/2020.01.21.914895

**Authors:** Jiaojiao Wang, Chuanxin Wu, Yunying Wang, Chongxiang Chen, Jing Cheng, Xiaolong Rao, Hang Sun

## Abstract

**Background:** High mobility group box 1 (HMGB1) is an important “late” inflammatory mediator in bacterial sepsis. Ethyl pyruvate (EP), an inhibitor of HMGB1, can prevent bacterial sepsis by decreasing HMGB1 levels. However, the role of HMGB1 in fungal sepsis is still unclear. Therefore, we investigated the role of HMGB1 and EP in invasive *C. albicans* infection.

**Methods:** We measured serum HMGB1 levels in patients with sepsis with *C. albicans* infection and without fungal infection, and control subjects. We collected clinical indices to estimate correlations between HMGB1 levels and disease severity. Furthermore, we experimentally stimulated mice with *C. albicans* and *C. albicans* + EP. Then, we examined HMGB1 levels from serum and tissue, investigated serum levels of tumor necrosis factor α (TNF-α) and interleukin 6 (IL-6), determined pathological changes in tissues, and assessed mortality.

**Results:** Serum HMGB1 levels in patients with severe sepsis with *C. albicans* infection were elevated. Increased HMGB1 levels were correlated with procalcitonin (PCT), C-reactive protein (CRP), 1,3-β-D-Glucan (BDG) and *C. albicans* sepsis severity. HMGB1 levels in serum and tissues were significantly increased within seven days after mice were infected with *C. albicans*. The administration of EP inhibited HMGB1 levels, decreased tissue damage, increased survival rates and inhibited the release of TNF-α and IL-6.

**Conclusions:** HMGB1 levels were significantly increased in invasive *C. albicans* infections. EP prevented *C. albicans* lethality by decreasing HMGB1 expression and release. HMGB1 may provide an effective diagnostic and therapeutic target for invasive *C. albicans* infections.

## **1.** Introduction

The incidence of invasive fungal infection (IFI), more than 50% of which is caused by *C. albicans*, has been steadily increasing and is the fourth most common cause of nosocomial bloodstream infection. This is in part due to invasive surgical operations, chemotherapy, radiotherapy, immunosuppressive therapy, and the wide use of broad-spectrum antibiotics and glucocorticoids(1–5). IFI is a serious infectious disease with a high mortality rate; its diagnosis and treatment are difficult in the clinical setting(6–8). Studies have indicated that proinflammatory cytokines, such as tumor necrosis factor α (TNF-α), interleukin 6 (IL-6) and high mobility group box 1 (HMGB1) influence tissue damage, hypotension, multiple organ failure and death in IFI(9–13). Therefore, the targeted therapy of these proinflammatory cytokines may improve the prognosis of IFI.

Previous studies have indicated that early proinflammatory cytokines release within one to two hours of sepsis onset, and peak at 24 hours(14). One reason for the ineffective performance of early proinflammatory cytokine antagonist therapies in clinical trials is that when sepsis is diagnosed, proinflammatory cytokines have already been released in large quantities, and therefore, the optimal treatment time is lost. Recently, HMGB1 has been recognized as a key “late” inflammatory mediator in lipopolysaccharide (LPS)-induced sepsis(15). HMGB1 is released at approximately 12–18 h after macrophages with LPS stimulation and then is maintained at high levels. Similar HMGB1 delayed kinetics are also observed in mice serum during lethal endotoxemia. The administration of HMGB1 inhibitors, such as Anti-HMGB1 mAb, before or after lethal endotoxemia, prevents endotoxin-induced death, inhibiting the release of HMGB1, TNF-α and IL-6, improving survival rates and limiting tissue damage(15). Taken together, HMGB1 is a vital “late” inflammatory mediator of bacterial sepsis, and its delayed kinetics may provide a broader therapeutic window for bacterial sepsis.

Ethyl pyruvate (EP), a stable lipophilic pyruvate derivative, has recently been discovered as a highly-effective and security HMGB1 inhibitor. It has anti-inflammatory and anti-oxidant properties, which protect cells from inflammatory injury both *in vivo* and *in vitro*, such as hemorrhagic shock, sepsis and ischemia-reperfusion injury(16–18). Ulloa *et al.* reported that EP inhibited HMGB1 expression and improved survival in mice with established endotoxemia or sepsis(18). These results indicate that EP is an effective therapeutic against bacterial sepsis, mainly via HMGB1 inhibition.

To date, HMGB1 research has mainly focused on bacterial sepsis, while the role of HMGB1 in fungal sepsis is unknown. We speculate that HMGB1’s role in bacterial sepsis is similar to that in fungal sepsis, as both pathogenic mechanisms depend on the activation of early and late cytokine responses. Here, we measured HMGB1 mRNA and protein levels in patients with severe sepsis with *C. albicans* infection, to evaluate the relationship between HMGB1 and disease severity. In addition, mice were stimulated with *C. albicans* and *C. albicans* combined with EP to observe HMGB1 mRNA and protein levels in serum and tissue, at different time points. We explored the role of HMGB1 in invasive *C. albicans* infection and aimed to provide an effective diagnostic and therapeutic target for invasive *C. albicans* infections.

## 2. Materials and methods

### 2.1 Human subjects

Our study included four patient populations: 23 control subjects (group 1), 35 patients with sepsis without fungal infection (group 2), 35 patients with severe sepsis without fungal infection (group 3), and 23 patients with severe sepsis with *C. albicans* infection (group 4). All patients were selected from the Intensive Care Unit (ICU) and the Department of Hepatobiliary Surgery, the Second Affiliated Hospital of Chongqing Medical University, China between May 2015 and May 2016. Patient selection was based on diagnostic criteria from the International Guidelines for Management of Severe Sepsis and Septic Shock 2012(19). Patients with *C. albicans* sepsis were accorded with diagnostic criteria for severe sepsis, and *C. albicans* cultures from peripheral bloods were positive at least once. Healthy participants were randomly selected from outpatients of the Second Affiliated Hospital of Chongqing Medical University, as controls. This study was conducted in accordance with the Declaration of Helsinki, and was approved by the Human Research Ethics Committee of Chongqing Medical University (CHICTR-OCC-13003185). Informed consent was obtained from participants, which were given a full explanation of the study.

### 2.2 Blood sample collection and assays

Peripheral blood samples were collected after an overnight fast, when the sepsis diagnosis was confirmed. Samples were collected in pyrogen-free tubes containing heparin (Bioendo, Xiamen, China). Approximately 250 μl peripheral blood was immediately added to 750 μl lysate solution (Bioteke, Beijing, China), and stored at -70°C. The remaining blood was centrifuged at 3500 rpm for 15 minutes, after which the serum was divided into five separate 1.5 ml tubes and stored at -70°C until assaying. Clinical indices, including blood routine, liver and kidney functions, coagulation metrics, C-reactive protein(CRP), procalcitonin(PCT) and 1,3-β-D-Glucan(BDG) were measured by the Department of Laboratory Medicine, the Second Affiliated Hospital of Chongqing Medical University.

### 2.3 Animal care

Male BALB/c mice (body weight: 20–25g and 6–8 weeks old) were obtained from the Experimental Animal Center of Chongqing Medical University. Mice were housed in a pathogen-free environment and given *ad libitum* access to food and water. Indoor temperature were maintained at 22 ± 2°C with an artificial light/dark period of 12 hours. All animal experimental operations were in accordance with Chongqing Management Approach of Laboratory Animal (chongqing government order NO.195).

### 2.4 Establishment of an immunosuppressive mouse model

Forty BALB/c male healthy mice were intraperitoneally injected with 100 mg/kg/d cyclophosphamide (Sigma, USA) for three days. Mouse blood was taken by tail-breaking at baseline (one day before cyclophosphamide injection) and on the third day of cyclophosphamide injection. Blood routines were measured by the Department of Laboratory Medicine, the Second Affiliated Hospital of Chongqing Medical University.

### 2.5 **Selection of optimum *C. albicans* cell densities**

*C. albicans* (ATCC 14053) was supplied by the microbiology laboratory at the Second Affiliated Hospital of Chongqing Medical University. Cells were cultured for 48 h in Sabao weak medium (Microbial reagent company, Hangzhou, China) at 37°C and serially passaged three times prior to infection. Different *C. albicans* inoculum densities were prepared using sterilized saline.

On the fourth day after immunosuppression, mice in each group (n = 8) were assigned different *C. albicans* cell density inoculums (1×10^5^, 2×10^5^, 5×10^5^, 1×10^6^, 1×10^7^ cells per ml). Mice were injected with 200 μl of these densities via the lateral tail vein(20). Mortality in each group was observed for 10 days after infection. Based on these mortality rates, the *C. albicans* inoculum 2×10^5^ was selected for subsequent experiments.

### 2.6 Establishment of an invasive *C. albicans* infection model in mice

Mice in the invasive *C. albicans* infection group (n = 12) were injected with 200 μl of the 2×10^5^ inoculum via the tail vein on the fourth day after immunosuppression. Mice in the immunosuppressive group (n = 3) were injected with 200 μl saline. Infected mice on days 1, 3, 5, and 7 after infection (n = 3 mice/time point) and mice in the immunosuppressive group (n = 3) were then euthanized and the lungs, liver and kidneys were aseptically removed. Part of the liver, lungs and kidney were weighed, homogenized in phosphate buffered solution (PBS) (8.06 mM sodium phosphate, 1.94 mM potassium phosphate, 2.7 mM KCl, and 137 mM NaCl) (pH 7.4), serially diluted, and plated in duplicate on Sabao weak medium and CHROMagar *Candida* medium (Microbial reagent company, Hangzhou, China). Plates were incubated at 37°C and *C. albicans* growth was assessed at 48 h. Part of the liver, lung and kidney were fixed in 4% paraformaldehyde for 24 h, after which tissues were dehydrated by conventional alcohol gradient processing, embedded in paraffin, serial sectioned and stained with hematoxylin-eosin (H&E) or periodic acid-Schiff (PAS).

### 2.7 EP treatment in mice infected with invasive *C. albicans*

EP (28 mM) (Sigma, St. Louis, MO) was prepared in solution with sodium (130 mM), potassium (4 mM), calcium (2.7 mM) and chloride (139 mM) (pH 7.0).

Eighty mice were randomly divided into four groups (n = 20 for each group): 1) the control group, 2) the immunosuppressive group, 3) the invasive *C. albicans* infection group, and 4) the EP group. Mice in the EP and the invasive *C. albicans* infection groups were intraperitoneally injected with 100 mg/kg/d cyclophosphamide for three days. On the fourth day, both groups were injected with 200 μl of the 2×10^5^ inoculum via the tail vein.

Mice in the EP group were intraperitoneally injected with 60 mg/kg EP four hours after infection, twice a day for three days. Mice in the invasive *C. albicans* infection group were injected with the same volume of saline. Three days after intraperitoneal injection of cyclophosphamide, the immunosuppressive group were given the same volume of saline. The control group were injected with the same volume of saline. In addition, the mortality in each group (n = 15) was observed during all procedures.

Clinical symptoms (e.g., appetite, activities and weight changes, etc.) were observed and recorded at 8:00 a.m. everyday. Mice were euthanized on days 1, 3, 5, and 7 after infection (n = 5 mice/time point). Blood was aseptically drawn from the orbital venous plexus and collected in pyrogen-free tubes containing heparin. Approximately 250 μl of peripheral blood was immediately added to 750 μl lysate solution (Bioteke, Beijing, China) and stored at -70°C. The remaining blood was centrifuged at 3500 rpm for 15 minutes at 4°C. The supernatant was collected in a clean micro-tube, which was stored at -70°C until assaying.

After euthanasia, the lungs, liver and kidneys were aseptically removed. Some sections were frozen in liquid nitrogen and other sections were fixed in 10% formalin and embedded in paraffin. Tissue sections were stained with H&E or placed on poly-L-lysine-coated glass slides for immunohistochemistry.

### 2.8 Quantitative real-time polymerase chain reaction (qRT-PCR) analysis

Total RNA was extracted using the total RNA isolation kit (Bioteke, Beijing, China), according to manufacturers’ instructions. The RNA was reverse transcribed into cDNA using the PrimeScript RT reagent kit (Takara, Dalian, China). Quantitative real-time PCR analysis was performed using the SYBR Premix Ex TaqTM II kit (Takara, Dalian, China). HMGB1 primers were designed using Primier4.1 software and GADPH was used as a reference gene. Primers were purchased from Huada, Dalian, China.

Human HMGB1 primers were: 5’-GCGGACAAGGCCCGTTA-3’ (sense) and 5’-AGAGGAAGAAGGCCGAAGGAA -3’ (antisense), and GADPH primers were 5’-TGCCAAATATGATGACATCAAGAA-3’ (sense) and 5’-GGAGTGGGTGTCGCTGTTG-3’ (antisense).

Mouse HMGB1 primers were: 5’-ACCCGGATGCTTCTGTCAACTTCT-3’ (sense) and 5’-GCCTTGTCAGCCTTTGCCATATCT-3’ (antisense); and mouse GADPH primers were 5’-CATGGCCTTCCGTGTTCCTA-3’ (sense) and 5’-GCGGCACGTCAGATCCA-3’ (antisense). The mRNA expression levels of target genes were normalized to GADPH using the 2^-ΔΔCt^ method, where ΔCt = target gene Ct - GAPDH Ct and ΔΔCt = ΔCt treatment-ΔCt control. Data values of 2^-ΔΔCt^ > 2, were considered statistically significant. Three independent experiments were performed in triplicate.

### 2.9 Enzyme-linked immunosorbent assay (ELISA) analysis

The recombinant HMGB1 (rHMGB1, Novoprotein, Shanghai, China) series concentrations, ranging from 15.625 ng/ml to 1 μg/ml, and serum diluted with PBS (1∶ 8) from patients or mice, were incubated overnight in 96-well plates at 4°C. Plates were then washed five times in PBS Tween-20 buffer (PBST; 0.05% Tween-20 in PBS). This step was performed after each incubation period. The plates were blocked for two hours at 37°C using 3% bovine serum albumin (BSA) in PBST. After this, rabbit anti-HMGB1 polyclonal antibody (1:1,500, Abcam, U.S.A), diluted in PBS buffer containing 2% skimmed milk, was added to all wells and incubated for two hours at 37°C. The plates were then incubated for one hour at the same temperature, with anti-IgG rabbit antibody conjugated to horseradish peroxidase (1:5,000, Golden Bridge Biotech Co Ltd, Beijing, China). After washing, tetramethylbenzidine (Abcam, U.S.A) was added to each well and plates were incubated in the dark for 10–20 minutes at 37°C. Reactions were monitored by measuring the absorbance at 460 nm (Thermo, U.S.A). All assays were run in duplicate on each test plate.

TNF-α and IL-6 levels were assessed using commercial ELISA kits (4A Biotech Co., Ltd, Beijing, China).

### 2.10 Western blot analysis

Tissue total proteins were extracted using the total protein isolation kit (Solarbio, Beijing, China), according to manufacturers’ instructions. Proteins were quantified using the Nanodrop 1000 (Thermo Scientific, Wilmington, Del.). Samples were denatured at 100°C for five minutes. Equal protein concentrations were loaded onto 12% SDS-PAGE gels and transferred to polyvinylidene fluoride membranes (Millipore Corporation, Billerica, Mass.). Membranes containing proteins were blocked in 5% BSA and incubated with rabbit anti-HMGB1 polyclonal antibody (1:1,000, Abcam, U.S.A), followed by incubation with horseradish peroxidase-linked secondary antibodies (1:5,000; Golden Bridge Biotech Co. Ltd, Beijing, China). For standardization and expression comparisons, the membranes were also hybridized with a primary anti-β-actin antibody (1:1,000; Wuhan Boster Biological Technology, Wuhan, China). The bands appearing on film were analyzed with GeneTools software (Syngene, Frederick, Md).

### 2.11 Immunohistochemistry

Tissue sections on poly-L-lysine-coated glass slides were deparaffinized in xylene and rehydrated in a graded alcohol series. Then the slides were blocked by 3% hydrogen peroxide for 10 minutes at room temperature (RT), and antigen retrieval for 15 minutes at 100°C. The slides were then incubated in 5% BSA for one hour to block nonspecific protein binding sites. After washing in PBS, the slides were incubated with an anti-HMGB1 polyclonal antibody (1:1,000; Abcam, U.S.A) overnight at 4°C. PBS, without primary antibody, was used as a negative control. The slides were washed three times in PBS, then incubated with horseradish peroxidase-linked secondary antibodies (1:5,000; Golden Bridge Biotech Co. Ltd) for 15–20 minutes at RT. Slides were colored with diaminobenzidine and hydrogen peroxide (Golden Bridge Biotech Co Ltd), counterstained with hematoxylin, dehydrated and mounted for viewing (Nikon, Japan). Two different sections from each organ, per time point, were assessed per experiment.

### 2.12 Statistical analysis

We used SPSS version 20.0 software for statistical analysis. Data were displayed as the mean ± SD. Categorical variables were analyzed using χ2 or Fisher exact tests. Bonferroni tests were applied when normality and homogeneity of variance were satisfied, otherwise Tamhane’s T2 tests were applied. Differences in survival rates among groups were compared by the Log-Rank test. A *P*-value <0.05 was considered statistically significant.

## 3. Results

### 3.1 Demographics and clinical characteristics

Whole blood and serum from 116 participants (35 patients with sepsis without fungal infection, 35 patients with severe sepsis without fungal infection and 23 patients with severe sepsis and *C. albicans* infection and 23 healthy controls) were collected. The clinical characteristics of these subjects are detailed (Table 1). No significant differences in age and gender were observed between all groups.

**Table 1.** Demographics and clinical characteristics.

| Group |  | 1 | 2 | 3 | 4 |
| --- | --- | --- | --- | --- | --- |
| Gender (F/M) |  | 10/13 | 12/23 | 11/24 | 11/12 |
| Age (y) |  | 62.2±16.3 | 60.5±17.1 | 62.8±15.5 | 65.2±23.8 |
| Blood routine | WBC (×10<sup>9</sup>/L) | 6.7±3.2** | 13.3±4.2* | 15.1±7.7 | 15.9±5.1 |
|  | NEUT (%) | 55.4±2.9** | 86.1±5.7 | 92.2±4.2* | 82.8±11.4 |
|  | RBC (×10<sup>12</sup>/L) | 4.3±0.5** | 3.6±0.5 | 3.8±0.3* | 3.2±0.7 |
|  | HGB (g/L) | 133.5±8.2* | 105.5±16.9 | 112.7±9.9 | 96.8±22.4 |
|  | PLT (×10<sup>9</sup>/L) | 274.3±50.4 | 246.2±112.8 | 138.5±74.5 | 167.8±77.0 |
| Liver function | TP (g/L) | 70.1±5.5 | 66.0±6.0 | 54.7±7.6* | 65.0±9.4 |
|  | ALB (g/L) | 47.1±7.9* | 32.8±4.5 | 29.6±4.9 | 29.0±4.7 |
|  | ALT (U/L) | 21.5±14.2 | 34.8±27.9 | 155.6±186.2 | 28.5±14.9 |
|  | AST (U/L) | 15.9±4.6** | 40.8±37.6 | 171.7±165.8 | 61.3±44.9 |
|  | ALP (U/L) | 72.8±29.9 | 77.3±20.8** | 160.5±178.6 | 243.8±174.7 |
|  | GGT (U/L) | 23.5±9.1** | 40.6±18.9* | 284.7±343.2 | 228.8±242.9 |
|  | TB (umol/L) | 11.6±2.1 | 10.64±4.3 | 58.7±52.1 | 38.6±44.1 |
|  | DB (umol/L) | 7.4±1.5* | 5.2±1.4* | 38.3±31.1 | 32.2±41.4 |
| Renal function | BUN (mmol/L) | 4.9±1.5** | 5.3±2.2** | 11.2±8.2 | 15.0±8.6 |
|  | SCr (umol/L) | 58.4±19.2 | 63.8±17.4 | 167.5±159.1 | 118.8±98.1 |
| Blood coagulation | PTA (%) | 94.1±11.8* | 82.4±12.9 | 60.2±17.5 | 63.7±23.7 |
|  | PT (S) | 12.3±1.0* | 14.7±1.5 | 17.9±2.7 | 16.8±2.5 |
|  | APTT (S) | 38.4±6.7 | 40.2±3.2 | 46.2±12.4 | 51.2±11.8 |
|  | FIB (g/L) | 3.4±0.5 | 6.0±2.2 | 4.8±1.3 | 4.8±2.0 |
|  | TT (S) | 17.9±0.9 | 16.1±1.2 | 16.5±1.3 | 17.3±0.4 |
| PCT (ng/ml) |  | 0.4±0.1** | 10.8±24.6 | 97.5±222.5 | 3.5±2.1 |
| CRP (mg/L) |  | 7.3±0.4** | 75.7±51.9 | 150.8±67.3 | 106.9±21.8 |
| BDG (pg/ml) |  | 50.2±21.0** | 61.5±24.7** | 61.5±24.7** | 268.9±88.4 |

Notes: group 1, control subjects; group 2, patients with sepsis without fungal infection; group 3, patients with severe sepsis without fungal infection; group 4, patients with severe sepsis with *C. albicans* infection. \*: $P < 0.05$ , \*\*: $P < 0.01$ , compared with group 4. WBC, white blood cells; NEUT%, neutrophil percent; RBC, red blood cells; HGB, hemoglobin; PLT, platelets; TP, total protein; ALB, albumin; ALT, alanine aminotransferase; AST, aspartate aminotransferase; ALP, alkaline phosphatase; GGT, $\gamma$ -glutamyl transpeptidase; TB, total bilirubin; DB, direct bilirubin; BUN, blood urea nitrogen; SCr, serum creatinine; PTA, prothrombin activity; PT, prothrombin
time; APTT, activated partial thromboplastin time; FIB, fibrinogen; TT, thrombin time; PCT, procalcitonin; CRP, C-reactive protein.

### 3.2 HMGB1 mRNA levels in patients with severe sepsis and *C. albicans* infection

In comparison with healthy controls (group 1), HMGB1 mRNA levels in patients with severe sepsis and *C. albicans* infection (group 4), in patients with severe sepsis without fungal infection (group 3) and patients with sepsis without fungal infection (group 2) were 3.66 ± 0.40, 3.12 ± 0.18 and 2.68 ± 0.35, respectively (Figure 1A). The 2^-ΔΔCt^ HMGB1 values for all three sepsis groups were greater than 2, suggesting that when compared with healthy controls, HMGB1 mRNA levels in these groups were significantly increased (*P*<0.01). Additionally, HMGB1 mRNA levels in group 4 were significantly higher than both groups 2 and 3 (*P*<0.05 or *P*<0.01) and group 3 levels were significantly higher than group 2 (*P*<0.05). These observations indicated that increased HMGB1 mRNA levels were positively correlated with sepsis severity.

**Figure 1.**
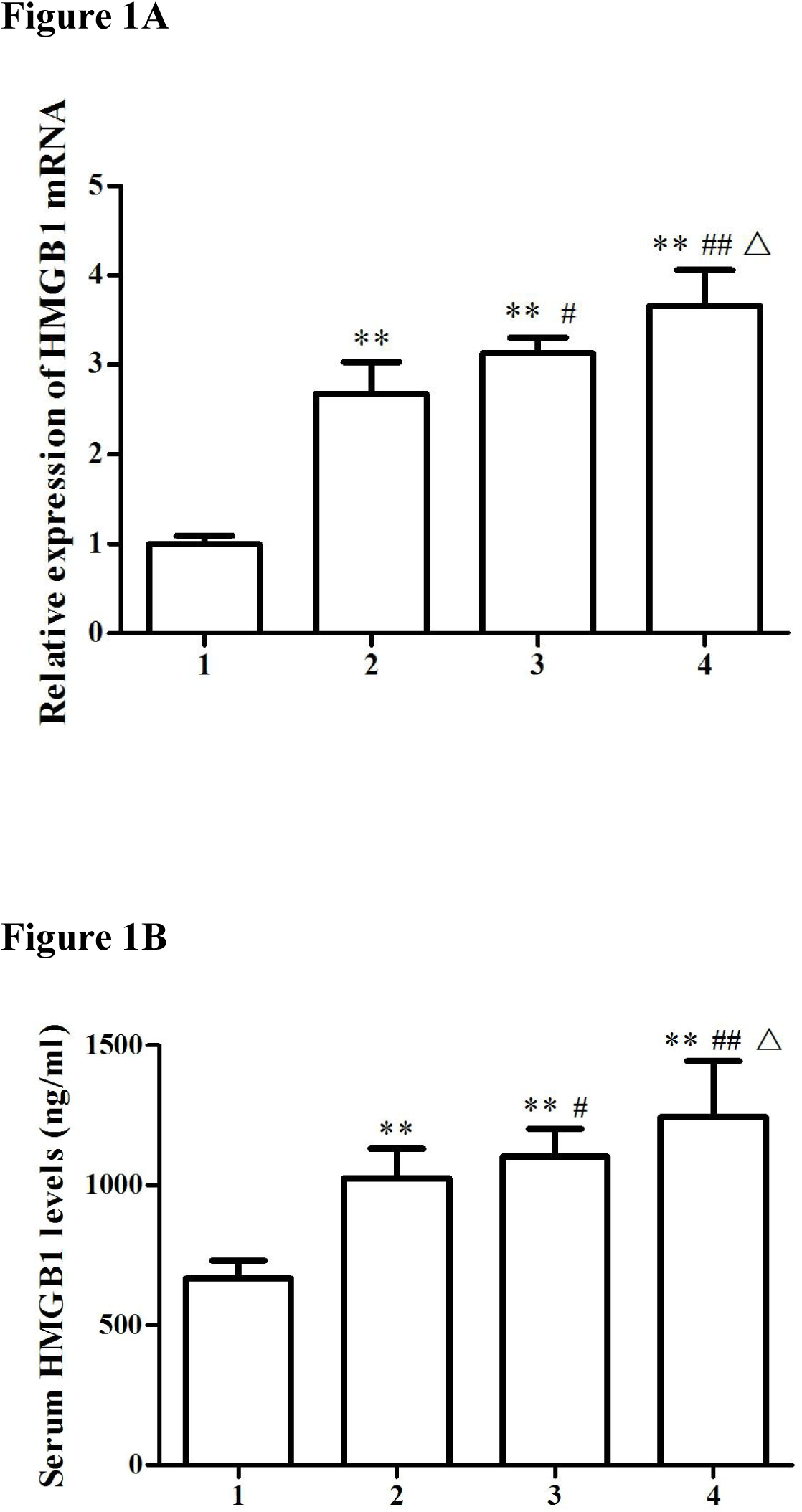
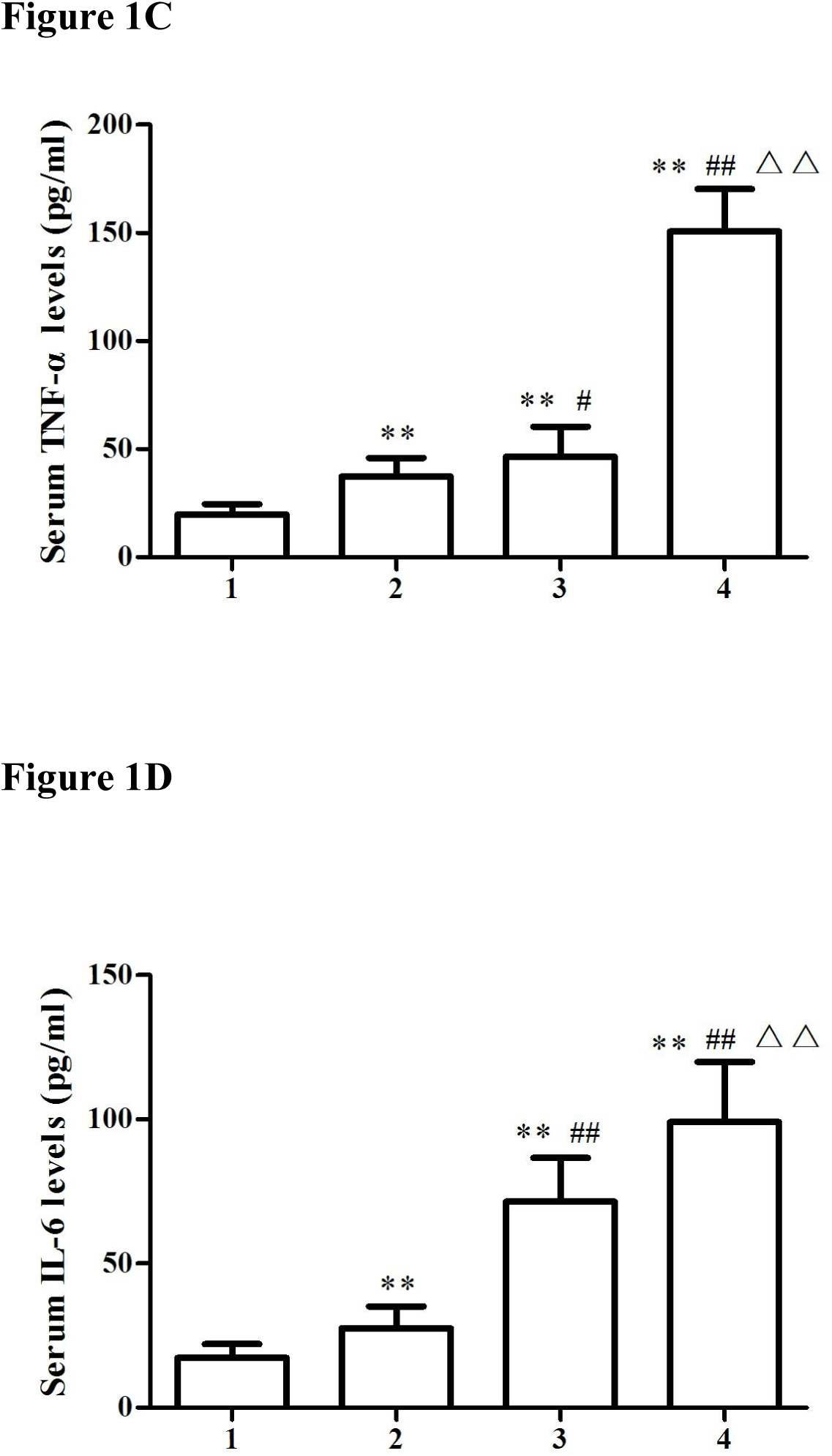
Expression of HMGB1, TNF-α and IL-6 in control subjects (group 1), sepsis patients without fungal infection (group 2), severe sepsis patients without fungal infection (group 3) and severe sepsis patients with *C. albicans* infection (group 4). A) HMGB1 mRNA expression was determined by RT-PCR. **: *P*<0.01, compared with group 1; #: *P*<0.05, ##: *P*<0.01, compared with group 2; Δ: *P*<0.05, compared with group 3. B–D) HMGB1, TNF-α and IL-6 serum levels were determined by ELISA. **: *P*<0.01, compared with group 1; #: *P*<0.05, ##: *P*<0.01, compared with group 2; Δ: *P*<0.05, ΔΔ: *P*<0.01, compared with group 3. Data are shown as the mean ± SD.

### 3.3 Serum HMGB1, TNF-α and IL-6 levels in patients with severe sepsis and *C. albicans* infection

Serum HMGB1 concentrations were 1243.83 ± 200.76 ng/ml in severe sepsis patients with *C. albicans* infection (group 4), 1101.67 ± 100.41 ng/ml in severe sepsis patients without fungal infection (group 3), 1025.94 ± 104.14 ng/ml in sepsis patients without fungal infection (group 2) and 666.44 ± 64.66 ng/ml in the control group (group 1). Statistical differences were observed across the four groups (*P*<0.05 or *P*<0.01). HMGB1 levels increased in all patients with sepsis versus the control group (*P*<0.01). Group 4 patients had statistically higher HMGB1 levels than groups 3 and 2 (*P*<0.05 and *P*<0.01) and group 3 patients had statistically higher HMGB1 levels than group 2 (*P*<0.05) (Figure 1B).

TNF-α and IL-6 levels in patient serum were, respectively; 150.74 ± 19.56 pg/ml and 99.06 ± 20.92 pg/ml in group 4, 46.49 ± 14.00 pg/ml and 71.38 ± 15.25 pg/ml in group 3, 37.34 ± 8.56 pg/ml and 27.59 ± 7.63 pg/ml in group 2, and 19.82 ± 4.82 pg/ml and 17.28 ± 4.75 pg/ml in the control group. All sepsis patients displayed statistically increased TNF-α and IL-6 levels when compared with the control group (*P*<0.01). There was evidence of significantly higher levels in group 4 when compared to groups 3 and 2 (*P*<0.01), and group 3 levels were significantly higher than group 2 (*P*<0.05 or *P*<0.01) (Figure 1C and 1D).

### 3.4 Correlations between HMGB1 and clinical indices

Correlation analyses revealed that HMGB1 serum levels in patients with severe sepsis and *C. albicans* infection were positively correlated with PCT, CRP and BDG (r = 0.491, 0.521, 0.504, *P*<0.05), respectively, but were not associated with WBC (r = 0.530, *P*>0.05) (Figure 2). HMGB1, PCT, CRP and BDG levels were increased with the development of *C. albicans* sepsis and were consistent with disease severity.

**Figure 2.**
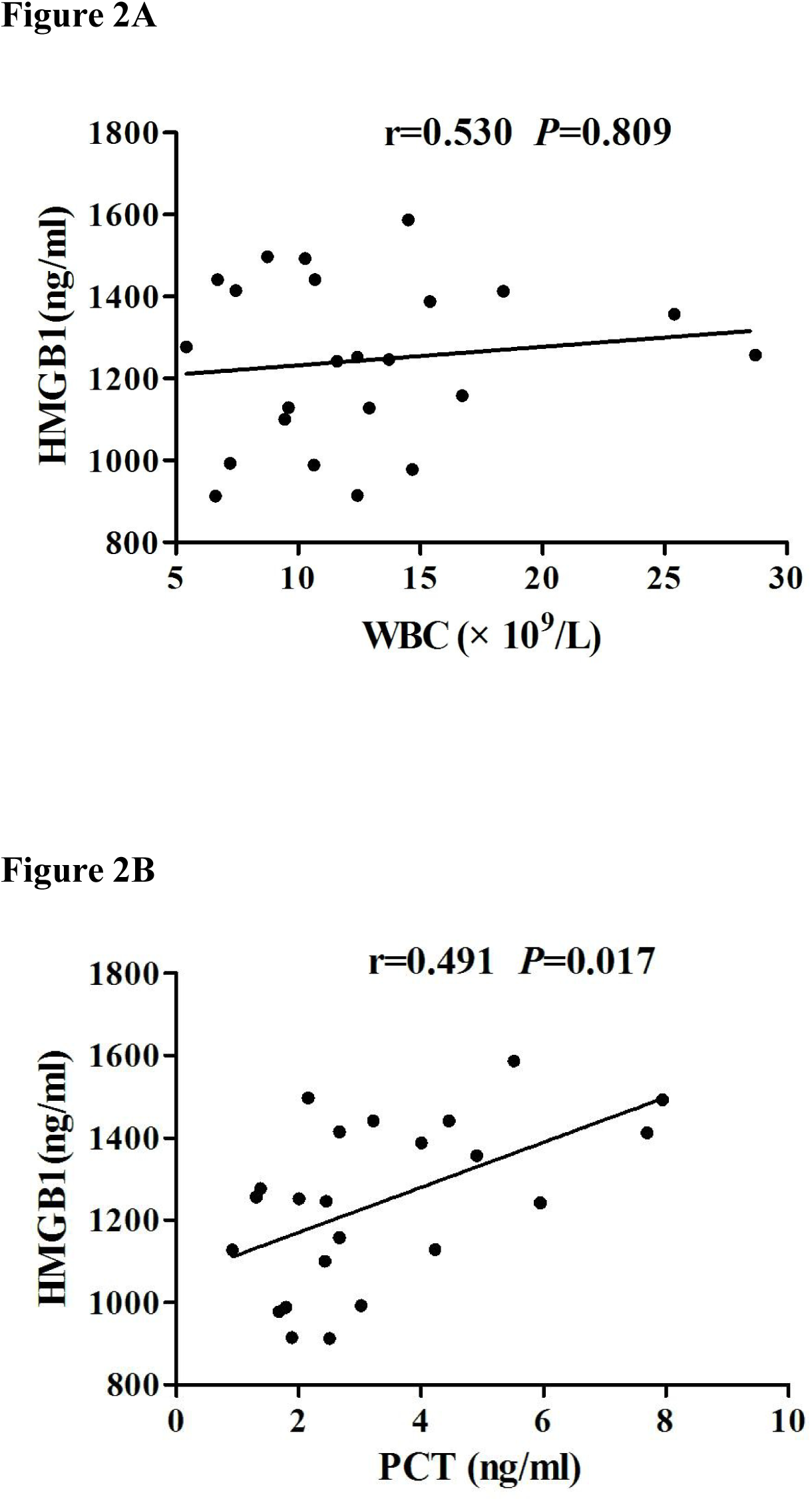
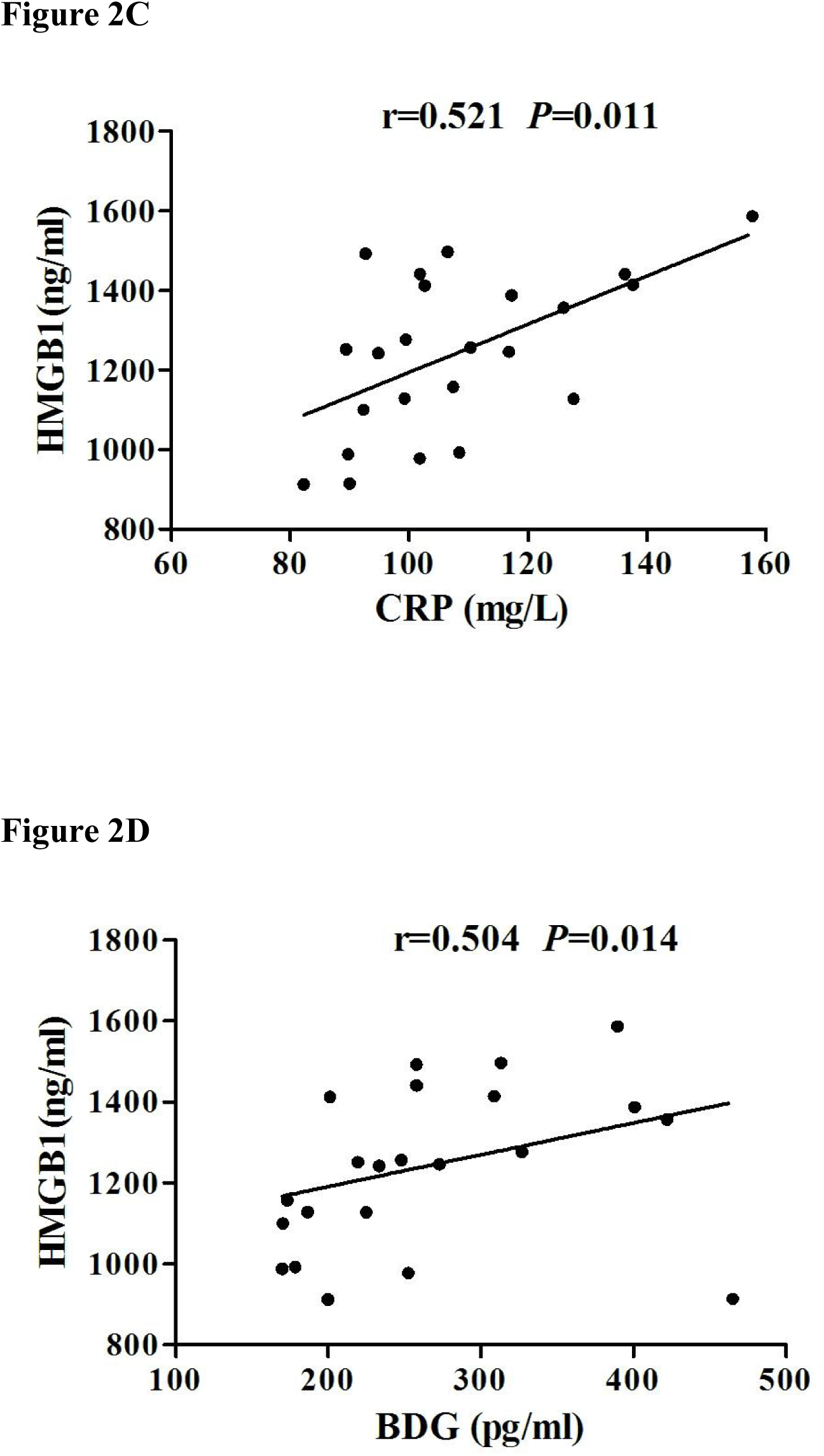
Correlation analysis between HMGB1 expression and WBC, PCT, CRP and BDG in patients with severe sepsis with *C. albicans* infection. A) The correlation between HMGB1 serum levels and WBCs; r = 0.530, *P* = 0.809. B) The correlation between HMGB1 serum levels and PCT; r = 0.530, *P* = 0.017. C) The correlation between HMGB1 serum levels and CRP; r = 0.521, *P* = 0.011. D) The correlation between HMGB1 serum levels and BDG; r = 0.504, *P* = 0.014.

### 3.5 Establishment of a mouse model for invasive *C. albicans* infections

After mice were injected with cyclophosphamide, routine blood analyses showed that WBCs, red blood cells (RBCs), hemoglobin (HGB) and platelets (PLTs) were significantly decreased, notably WBCs, indicating that the immunosuppressive mouse model was successfully established (Table 2). Mice (n = 8 per group) from different groups were injected with different *C. albicans* inoculums after immunosuppression, to investigate mortality rates. Based on these rates, the 2×10^5^ inoculum was selected for subsequent experiments (Figure 3A).

**Figure 3.**
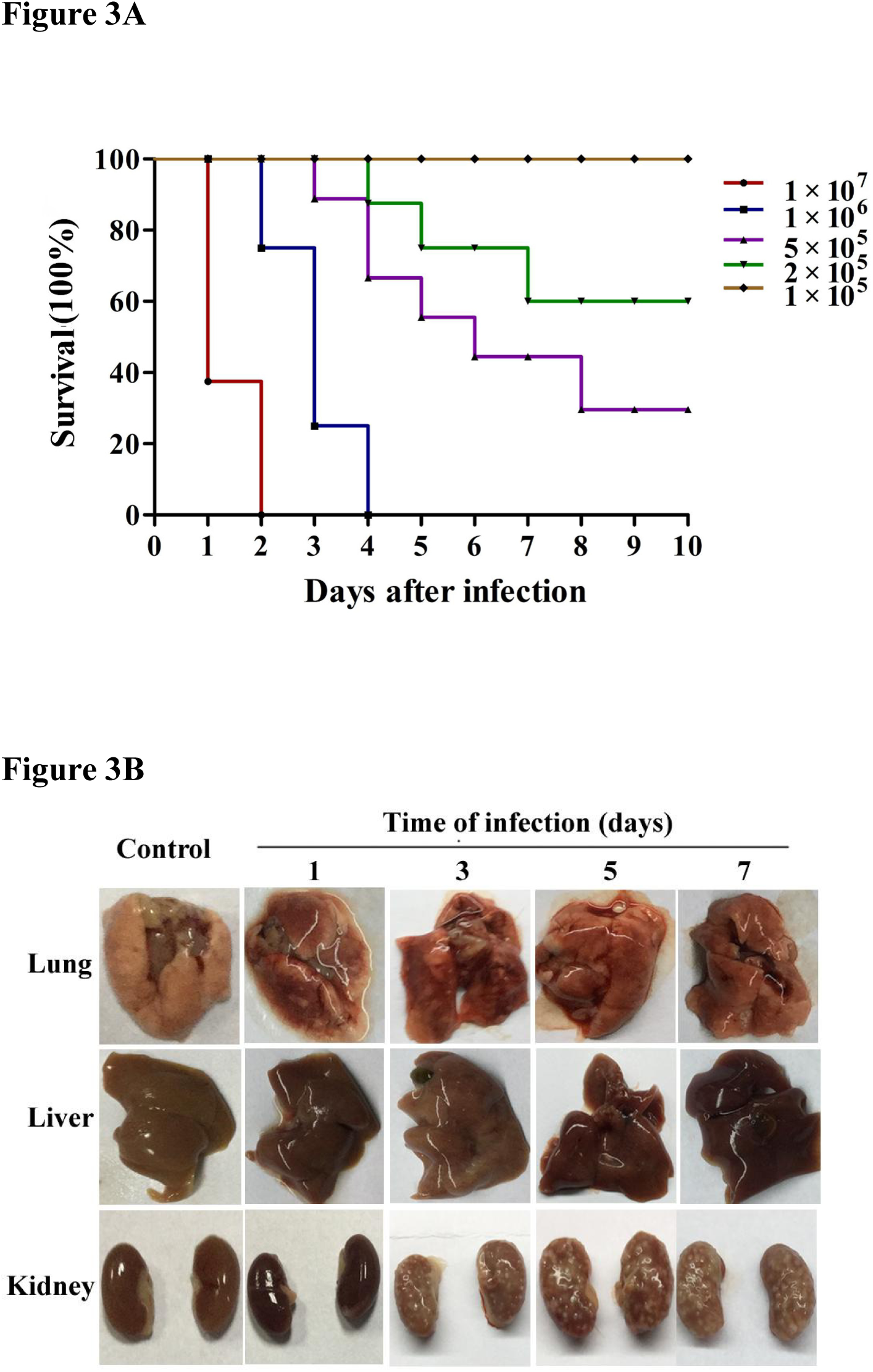
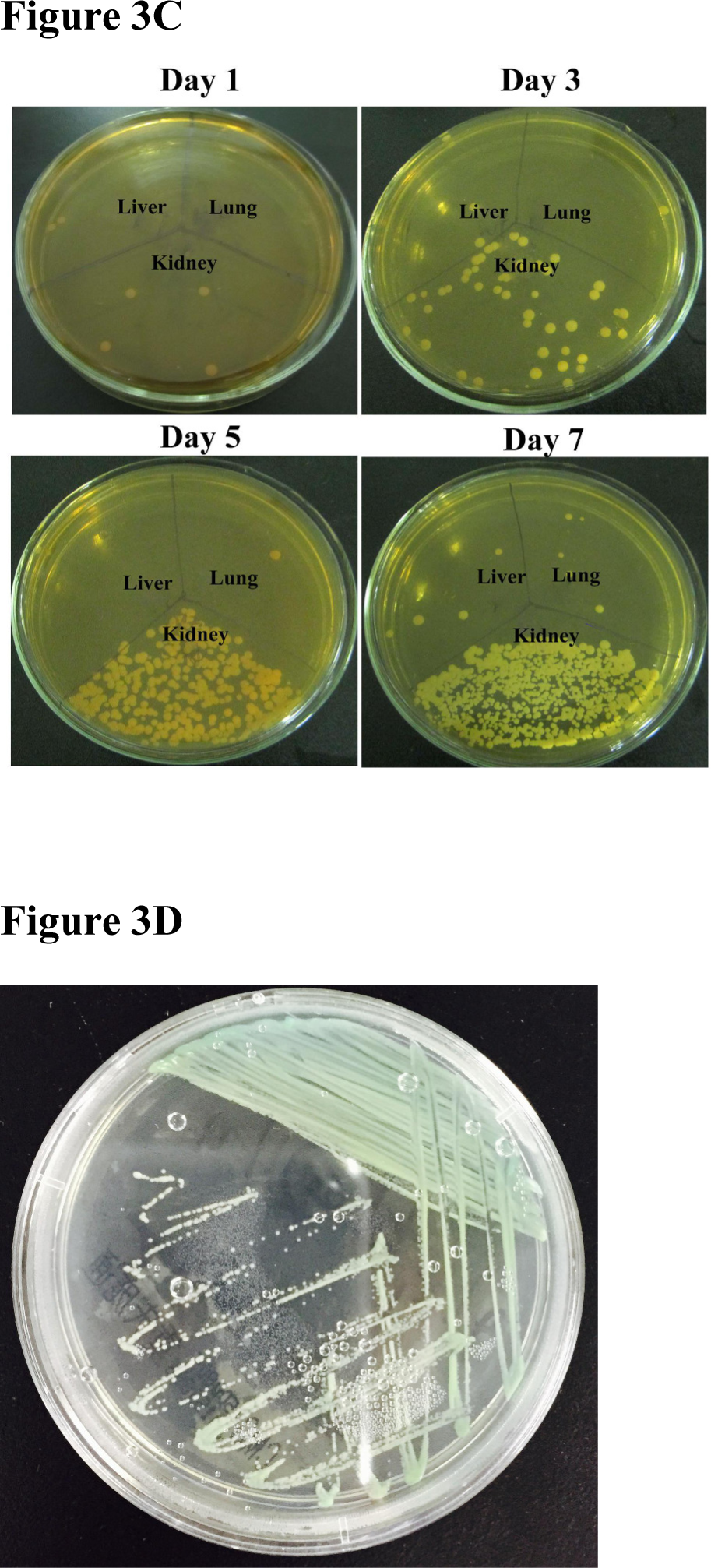
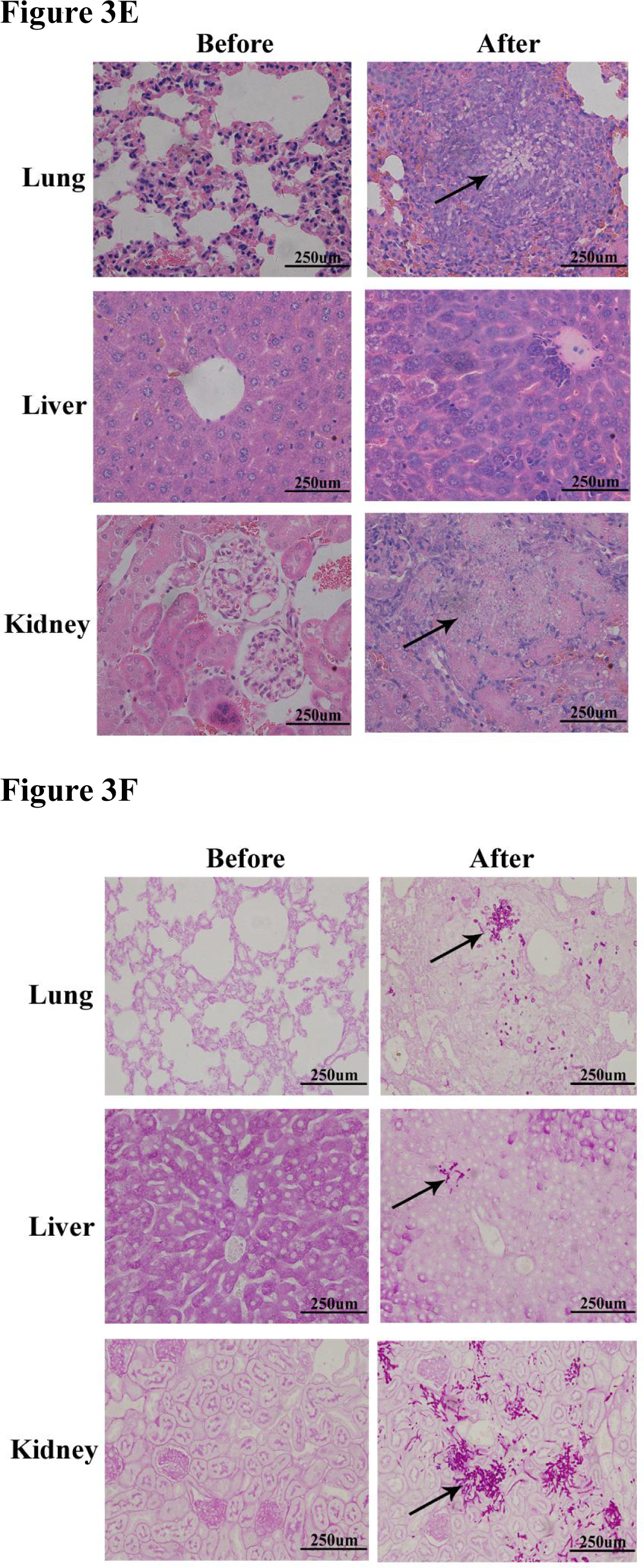
Establishment of a mouse model for invasive *C. albicans* infection. A) Survival curves of mice infected with different inoculums of *C. albicans*. According to survival curves, we selected a 2×10^5^ inoculum for subsequent experiments. B) Morphological characteristics of lung, liver and kidney before and after *C. albicans* infection. C) Fungal loads in lung, liver and kidney tissues after infection *C. albicans* growth was determined after a 48 hour incubation at 37°C on Sabao weak medium. D) Identification of *C. albicans* in lung, liver and kidney tissues using CHROMagar. E) H&E staining of lung, liver and kidney before and after *C. albicans* infection. Original magnification; ×400. Arrows show hyphae and spores. F) PAS staining of lung, liver and kidney before and after *C. albicans* infection. Magnification: ×400. Arrows show hyphae and spores.

**Table 2.** Hematocyte variables before and after cyclophosphamide administration.

|  | Before | After |
| --- | --- | --- |
| **WBC ($\times 10^9/\mu\text{l}$)** | 7.69±2.31 | 1.74±0.54** |
| **NEUT# ($\times 10^9/\mu\text{l}$)** | 0.17±0.09 | 0.13±0.20 |
| **LYMPH# ($\times 10^9/\mu\text{l}$)** | 7.04±2.08 | 1.35±0.45** |
| **MONO# ($\times 10^9/\mu\text{l}$)** | 0.11±0.10 | 0.02±0.02** |
| **RBC ($\times 10^{12}/\mu\text{l}$)** | 9.35±0.58 | 7.05±1.12** |
| **HGB (g/L)** | 140.82±8.30 | 106.56±16.30** |
| **PLT ($\times 10^9/\mu\text{l}$)** | 799.57±111.82 | 701.71±133.88* |

Notes: \*: $P < 0.05$ , \*\*: $P < 0.01$ , compared with before injection of cyclophosphamide. WBC, white blood cell; NEUT#, neutrophil count; LYMPH#, lymphocyte count; MONO#, monocyte count; RBC, red blood cell; HGB, hemoglobin; PLT, platelet.

After mice were infected with *C. albicans* and subsequent tissue removal, pulmonary tissue showed slight hyperemia, and white punctate necrosis could be seen on tissue surfaces on days one and three. The liver was slightly enlarged and hyperemic. From the third day after infection, white necrosis spots were seen (faintly) on the kidney surface, especially on days five and seven (Figure 3B).

It has been reported that fungal loads can vary in organs of mice with invasive *Candida* infection, with kidneys being consistently identified as the organ with the highest fungal load(20). In our experiments, a slight fungal presence was observed in the lung and liver after infection with *C. albicans,* but this was increased in the kidneys (Figure 3C). Fungi from these tissues were serially diluted and inoculated onto CHROMagar *Candida* medium where colonies grew green, indicating the presence of *C. albicans* (Figure 3D).

H&E and PAS staining demonstrated that *Candida* filamentous growth was observed in the kidney after infection, and mainly *Candida* spores were found in the lung and liver (Figure 3E–F). These data were consistent with previous reports(20), suggesting the successful constructed of a mouse model for invasive *C. albicans* infection.

### 3.6 Clinical manifestations and tissue damage in mice infected with *C. albicans*

After *C. albicans* infection, mice in the invasive *C. albicans* group suffered with diarrhea, decreased activity, lethargy, huddling, piloerection, weight loss, hepatorenal damage and death. Treatment with EP conferred significant protection from fatal *C. albicans* infection, which improved these clinical manifestations, such as reducing weight loss (Figure 4A), ameliorating the survival rate (mouse survival in the invasive *C. albicans* infection group = 10/15; mouse survival in the EP group = 13/15; *P*<0.05), (Figure 4B), and improving liver and kidney function (Table 3). Mouse activity, appetite and weight began to decrease early after cyclophosphamide injection and gradually improved at later time points (Figure 4A). No hepatorenal damage or death were found in the control and immunosuppressive groups during the experimental period.

**Figure 4.**
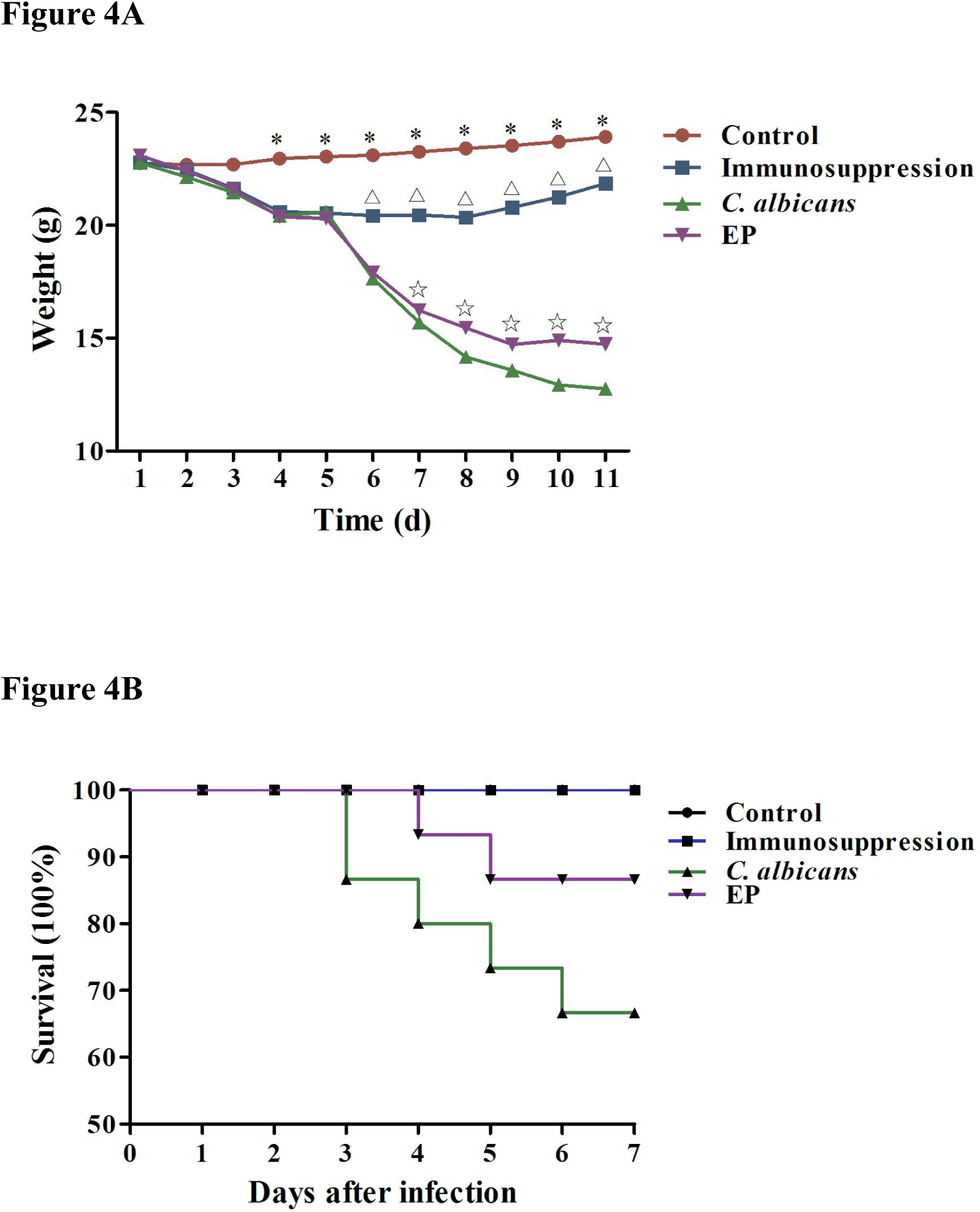
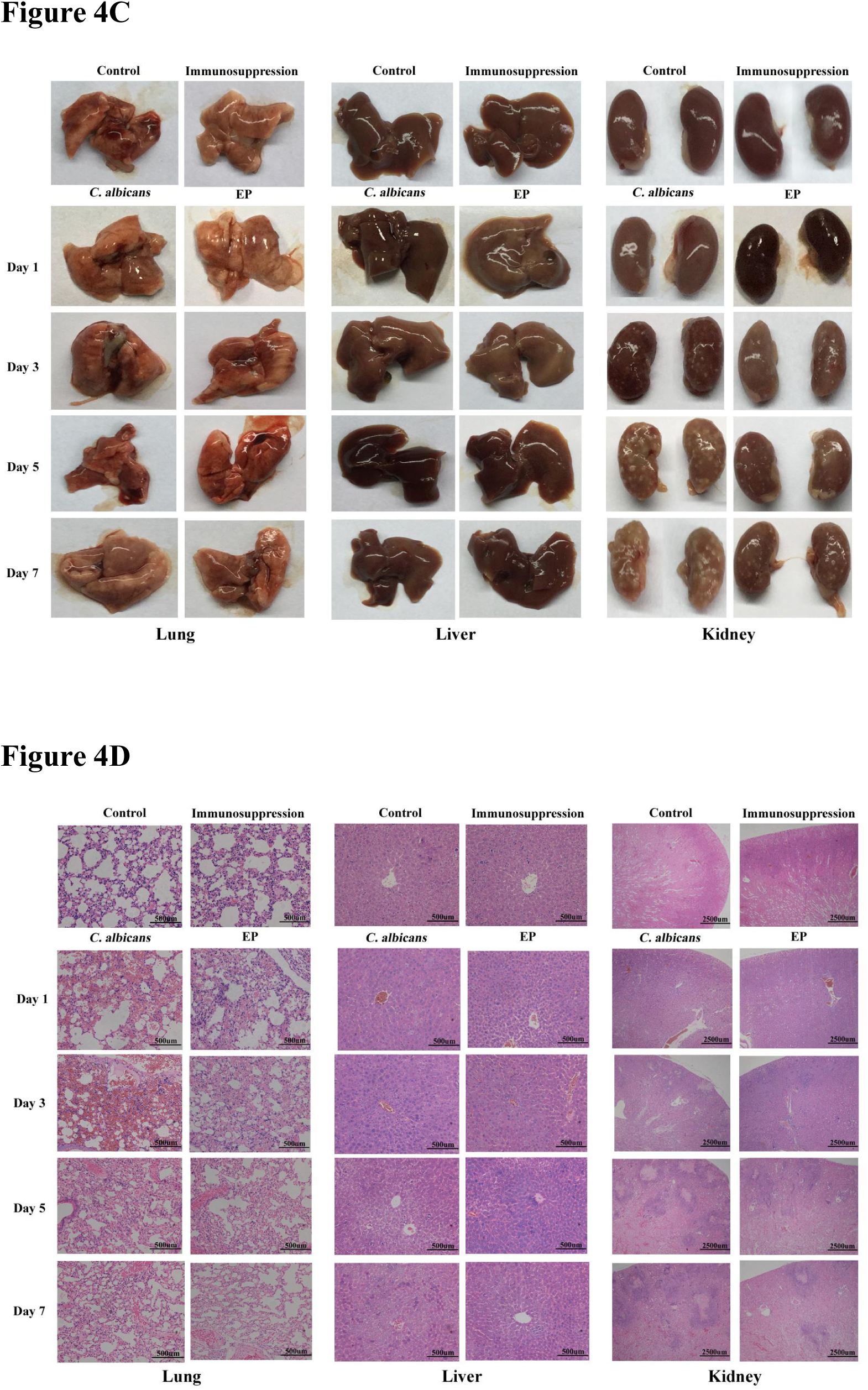
Clinical manifestations and tissue damage in mice infected with *C. albicans*. A) Changes in body weight of mice from control, immunosuppression, invasive *C. albicans* infection and EP groups. *: *P*<0.05 indicates differences from all other groups; Δ: *P*<0.05 indicates differences from *C. albicans* and EP groups; ✫: *P*<0.05 indicates differences from the EP group. B) The survival curve of controls, immunosuppression, invasive *C. albicans* infection and EP groups. C) Morphological characteristics of lung, liver and kidney of different mice groups at different times. D) H&E staining of lung, liver and kidney of different mice groups at different times. Magnification: ×200 (lung and liver), ×40 (kidney).

**Table 3.** Liver and kidney function in mice.

|  | Control | Immuno-suppression | *Candida albicans* |  |  |  | EP |  |  |  |
| --- | --- | --- | --- | --- | --- | --- | --- | --- | --- | --- |
|  |  |  | 1 | 3 | 5 | 7 | 1 | 3 | 5 | 7 |
| **TP(g/L)** | 52.1±2.6 | 52.2±3.6 | 50.2±3.1 | 70.7±1.5<sup>**</sup> | 76.6±3.5<sup>**</sup> | 87.1±3.1<sup>**</sup> | 49.6±1.3 | 61.5±2.9<sup>*△</sup> | 62.9±3.3<sup>*△△</sup> | 74.4±4.2<sup>**△</sup> |
| **ALB(g/L)** | 29.8±0.9 | 31.4±2.8 | 29.6±1.8 | 32.5±0.8 | 31.0±3.2 | 29.9±1.0 | 30.6±5.1 | 30.0±1.5 | 29.9±3.4 | 32.8±1.8 |
| **ALT(U/L)** | 78.3±6.0 | 76.3±6.2 | 68.1±8.5 | 137.3±12.7<sup>**</sup> | 158.7±13.2<sup>**</sup> | 164.0±5.1<sup>**</sup> | 66.9±6.3 | 107.4±11.3<sup>*△</sup> | 129.1±7.9<sup>**△</sup> | 144.0±8.5<sup>**△</sup> |
| **AST(U/L)** | 159.3±10.6 | 156.6±8.3 | 172.8±5.1 | 402.1±18.1<sup>**</sup> | 454.8±16.7<sup>**</sup> | 383.9±11.8<sup>**</sup> | 165.6±9.8 | 356.1±10.1<sup>**△</sup> | 390.1±30.6<sup>**△</sup> | 348.8±12.3<sup>**△</sup> |
| **BUN(mmol/L)** | 7.2±0.6 | 7.0±0.5 | 8.4±0.5 | 47.5±3.7<sup>**</sup> | 47.4±4.1<sup>**</sup> | 32.6±3.3<sup>**</sup> | 7.7±0.5 | 39.4±3.9<sup>**△</sup> | 38.1±4.7<sup>**△</sup> | 26.0±2.4<sup>**△</sup> |
| **SCr(ummol/L)** | 50.2±6.4 | 55.5±4.0 | 125.0±3.5<sup>**</sup> | 153.2±3.9<sup>**</sup> | 151.8±3.3<sup>**</sup> | 164.2±4.7<sup>**</sup> | 113.1±8.6<sup>**</sup> | 137.6±3.5<sup>**△</sup> | 137.9±4.3<sup>**△</sup> | 146.2±5.5<sup>**△</sup> |

Notes: \*: $P<0.05$ , \*\*: $P<0.01$ , compared with the control and immunosuppression group; △: $P<0.05$ , △△: $P<0.01$ , comparison of the EP group with the invasive *C. albicans* infection group. TP, total protein; ALB, albumin; ALT, alanine aminotransferase; AST, aspartate aminotransferase; BUN, blood urea nitrogen; SCr, serum creatinine.

Gross tissue specimens showed that after *C. albicans* infection, slight swellings and hyperemia were observed in lung and liver tissues. There were no significant differences between the EP group and the invasive *C. albicans* infection group. From the third day after infection, white necrosis spots on the kidney surface in the EP group were significantly reduced when compared to the invasive *C. albicans* infection group. This observation indicated that EP had a protective effect on kidney tissue damage caused by *C. albicans* infection. No pathological changes were observed in lung, liver and kidney tissues in control and immunosuppressive groups (Figure 4C).

Lung, liver and kidney histology were examined at different time points. The lung emerged capillary congestion, alveolar septum widening, and alveolar shrinkage after infection with *C. albicans*, especially on days one and three. Hepatocyte swelling and steatosis were observed and became aggravated as infection time progressed. A large number of necrotic foci were observed in the renal cortex on the third day after infection, which continued to increase during the infection course and extended to the medulla of the kidney on the fifth and seventh day after infection. This was accompanied by infiltration of *C. albicans* and inflammatory cells into necrotic foci. In addition, hemorrhaging was seen in the medulla of the kidney on the fifth and seventh day. Similar phenomena were observed in the lung, liver and kidney in the EP group, but the pathological manifestations were significantly reduced when compared to the invasive *C. albicans* infection group. H&E staining showed no significant differences in lung, liver and kidney between the control and the immunosuppressive group (Figure 4D).

### 3.7 HMGB1 expression at the mRNA and protein levels in mice infected with *C. albicans*

HMGB1 mRNA levels in peripheral blood, lung, liver and kidneys of the invasive *C. albicans* infection and EP groups were significantly increased after *C. albicans* infection (*P*<0.05 or *P*<0.01). When compared with the invasive *C. albicans* infection group, HMGB1 mRNA levels in the EP group began to decrease from day three after infection. The differences were significant (*P*< 0.01, Figure 5A). HMGB1 protein expression in the lung, liver and kidneys were markedly elevated after infection in the invasive *C. albicans* infection group and the EP group, but then decreased from day 1 after infection in the EP group when compared with the invasive *C. albicans* infection group (Figure 5B). There were no significant differences in HMGB1 mRNA and protein levels between the control group and the immunosuppressive group. These results suggested that EP prevented HMGB1 mRNA and protein expression, post *C. albicans* infection.

**Figure 5.**
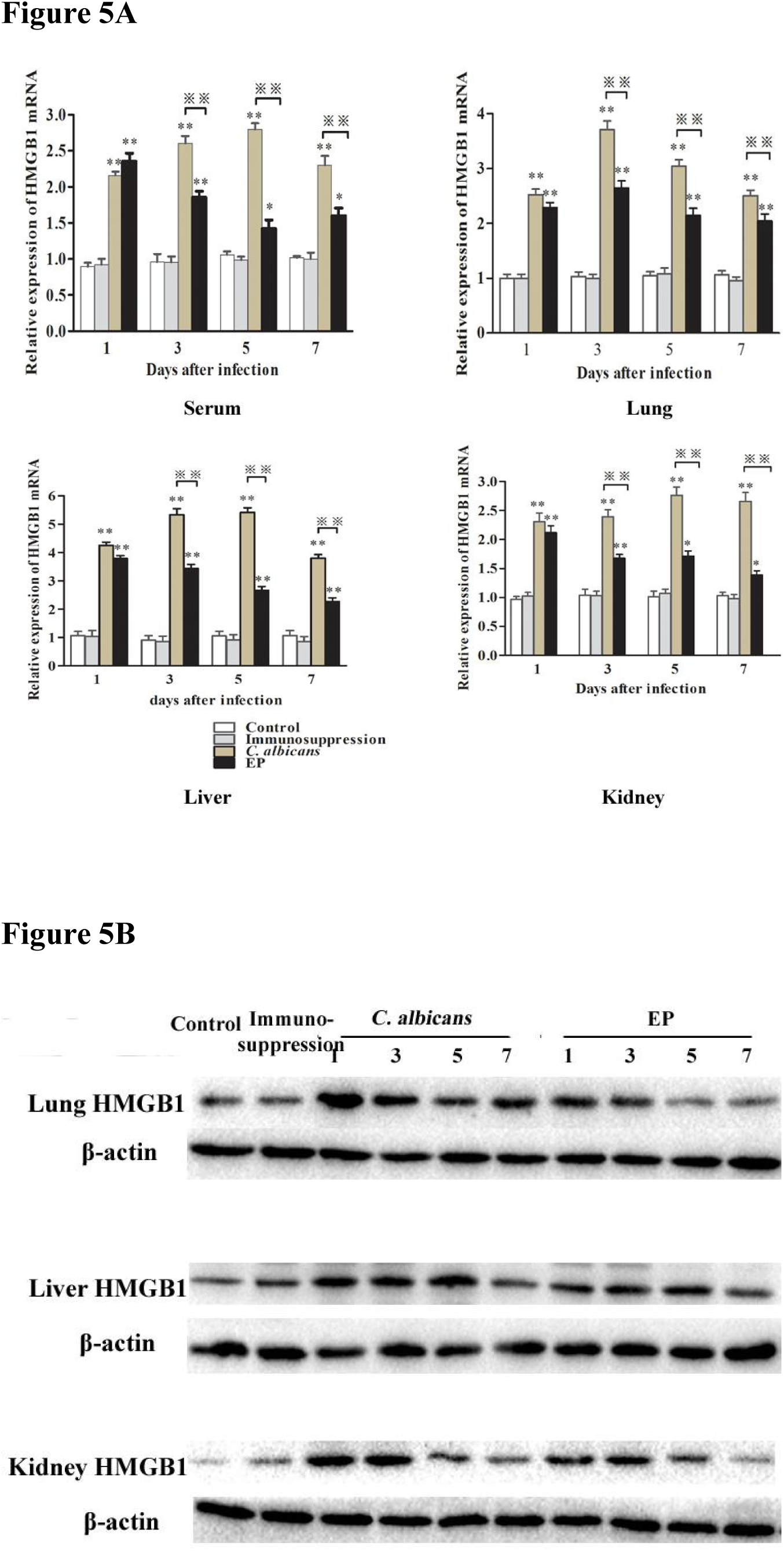
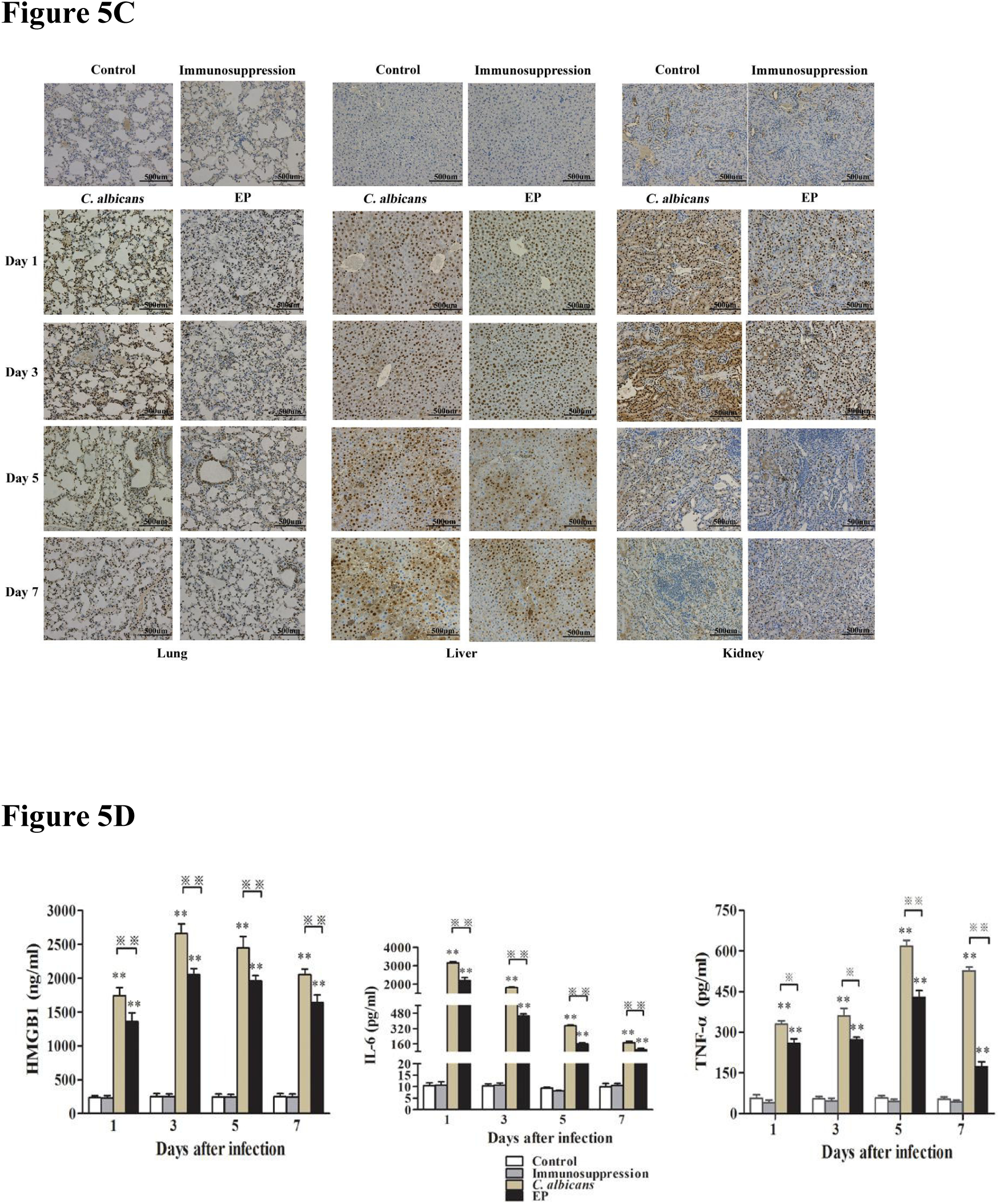
HMGB1 expression in mice groups. A) HMGB1 mRNA expression was determined by RT-PCR. Data are shown as the mean ± SD (n = 3). *: P<0.05, **: P<0.01, compare with control and immunosuppression group; ※※: P<0.01, compare EP group with invasive *C. albicans* infection group. B) Protein from lung, liver and kidney tissues and HMGB1 expression levels were detected by western blot. C) Immunohistochemical analysis of HMGB1 in lung, liver and kidney tissue. Magnification: ×200. D) HMGB1, TNF-α and IL-6 levels in mouse peripheral blood were evaluated by ELISA. Data are shown as the mean ± SD (n = 3). **: P<0.01, compare with control and immunosuppression groups; ※: P<0.05, ※※: P<0.01, compare EP group with invasive *C. albicans* infection group.

### 3.8 Relocation of HMGB1 in the lung, liver, and kidney of mice infected with *C. albicans*

In lung tissue of the control and immunosuppressive group, HMGB1 was expressed in the nucleus. There were no significant differences between the two groups. HMGB1 expression in the nuclei of the invasive *C. albicans* infection group and EP group increased significantly on the first, third, fifth and seventh day after infection, especially on days one and three. However, HMGB1 expression in the EP group declined when compared to the invasive *C. albicans* infection group at the same time point (Figure 5C).

In liver tissue, low HMGB1 expression was observed in the nuclei of the control and immunosuppressive group. On the first and third days after infection, nuclear HMGB1 expression in the invasive *C. albicans* infection group increased significantly, and low HMGB1 levels were released into the cytoplasm. On the fifth and seventh day after infection, in addition to increased HMGB1 expression in the nucleus, more HMGB1 was released into the cytoplasm. HMGB1 expression in the EP group was significantly lower when compared to the invasive *C. albicans* infection group at the same time point, suggesting that EP significantly inhibited HMGB1 release from the nucleus to cytoplasm (Figure 5C).

In kidney tissue, HMGB1 expression at the nucleus and cytoplasm increased significantly on day one and three after *C. albicans* infection and decreased gradually on days five and seven, but were still higher than the control and immunosuppressive group. This may have been due to the gradual increase of focal necrosis and severe tissue damage. HMGB1 expression in the EP group was significantly lower than the invasive *C. albicans* infection group at the same time point, indicating that EP not only inhibited HMGB1 expression in the nucleus, but also inhibited HMGB1 release from the nucleus to the cytoplasm (Figure 5C).

### 3.9 The release of inflammatory cytokines from the serum of mice infected with *C. albicans*

Serum HMGB1, TNF-α and IL-6 levels were significantly elevated on the first, third, fifth and seventh day after infection (*P*< 0.01), but were significantly decreased in the EP group when compared with the invasive *C. albicans* infection group, at the same time point (*P*<0.05 or *P*<0.01). There were no significant differences between the control and the immunosuppressive group. These results indicated that EP prevented the HMGB1 release after *C. albicans* infection (Figure 5D).

## 4. Discussion

Current diagnostic and treatment options for IFI are limited; therefore, it is essential to identify improved targets for diagnostic and therapeutic interventions. IFI is a systemic inflammatory response disease, where the overproduction of proinflammatory cytokines is believed to be a major cause of disease progression or death [9-13]. Of the numerous implicated inflammatory cytokines, HMGB1 appears to be a key mediator in IFI.

HMGB1 is an abundant chromatin-binding protein that mainly accumulates in the eukaryotic cell nucleus. It maintains nucleosome structure and is involved in the regulation of gene transcription(21, 22). Recently, it has been suggested that HMGB1 is released from immune cells under proinflammatory stimulus, and may be a vital late mediator in sepsis(23, 24). HMGB1 exerts its biological effects via receptor binding, including the receptor for advanced glycation end product (RAGE) and the Toll-like receptors. These interactions activate the nuclear factor-κB (NF-κB) and mitogen-activated protein kinase (MAPK) signaling pathways to release excessive levels of proinflammation cytokines, leading to increased inflammation(23, 24). Administration of HMGB1 inhibitors before or after lethal endotoxemia protects mice against lethal systemic inflammation, which may be an effective target for therapeutic intervention in bacterial sepsis. However, HMGB1 studies mainly focus on bacterial sepsis, with little on fungal sepsis.

In this study, HMGB1 mRNA and protein levels in patients of all three sepsis groups increased significantly when compared with healthy controls. Importantly, HMGB1 mRNA and protein levels in patients with severe sepsis with *C. albicans* infection were significantly higher than in patients with sepsis or severe sepsis without fungal infection. Also, in patients with severe sepsis without fungal infection, these levels were significantly higher than in patients with sepsis without fungal infection. These observations suggest that HMGB1 expression is significantly increased during sepsis, and these levels are positively correlated with sepsis severity.

Previous studies have also reported that HMGB1 expression reflects the severity of some infectious diseases, such as acute appendicitis and acute obstructive suppurative cholangitis, and helps clinicians estimate disease severity(25–27). In patients with *C. albicans* sepsis, monocytes/macrophages identify mannose molecules on cell walls, leading to increased proinflammatory cytokine release into the extracellular environment, including early cytokines, e.g., TNF, IL-1, IL-6 and HMGB1. Extracellular HMGB1 then acts as a proinflammatory mediator to induce multiple proinflammatory cytokine release, leading to inflammation and organ injury(12, 28). From this perspective, increased HMGB1 appears highly correlated with *C. albicans* sepsis severity, and may play an important role in the pathological progress of *C. albicans* sepsis. Controlling HMGB1 expression using anti-HMGB1 agents may improve the therapeutic treatment of *C. albicans* sepsis by managing systemic inflammation and reducing associated tissue injury.

WBCs, CRP and PCT are common inflammatory markers in clinical practice. In our study, serum HMGB1 levels in patients with severe sepsis with *C. albicans* infection were positively correlated with PCT, CRP and BDG, but not WBCs. WBCs may increase or decrease after infection, but they do not reflect infection severity(29). But, HMGB1 levels were correlated with *C. albicans* sepsis severity. Therefore, there are no correlations between HMGB1 and WBCs. Serum PCT and CRP levels are extremely low under normal circumstances, but increase significantly after sepsis, and increased PCT and CRP levels are typically correlated with sepsis severity(30, 31). However, PCT and CRP are limited in distinguishing between fungal infections and other pathogen infections. BDG is a cell wall component of most fungi, except for Zygomycetes and Cryptococcus species, and is important for the early diagnosis of IFI(32, 33). A recent meta-analysis showed that pooled sensitivity of BDG assay for the diagnosis of IFI was 76.8% [95% confidence interval (CI), 67–84%], and pooled specificity was 85.3% (95% CI, 80–90%)(34). BDG levels gradually decreased if patients responded to antifungal therapy(35). In our study, HMGB1, PCT, CRP and BDG levels were increased with the development of *C. albicans* sepsis and were consistent with disease severity, suggesting that HMGB1 may be a useful diagnostic and prognostic biomarker of *C. albicans* sepsis. Taken together, our data suggests that the combinatorial use of these indices may contribute to the early diagnosis of *C. albicans* sepsis and disease severity.

EP, a stable lipophilic pyruvate derivative, is a relatively nontoxic food additive as well as an HMGB1 inhibitor. After macrophages are stimulated with LPS, EP inhibits the release of HMGB1 and TNF-α and attenuates the activation of both the NF-κB and MAPK signaling pathways critical for cytokine release(18). Pretreatment or delayed administration of EP confers significant protection from lethal sepsis, reducing mortality by decreasing HMGB1 release(18). In our study, HMGB1 mRNA and protein levels in peripheral blood, lung, liver and kidney were significantly increased seven days after *C. albicans* infection, but were lower in the EP group than in the invasive *C. albicans* infection group, indicating that EP inhibited HMGB1 expression and release.

Previous studies have shown that HMGB1 release is the result of HMGB1 translocation from the nucleus to the cytoplasm(36, 37). HMGB1 released into the extracellular milieu may originate from prior synthetic reserves, while later new HMGB1 is synthesized in the nucleus and transferred to the cytoplasm, resulting in increased HMGB1 release. Released HMGB1, as a mediator of inflammatory disease, has the potential to be an ideal inflammatory biomarker and interventional target for therapy(38). In this study, nuclear HMGB1 in the liver was increased during the course of *C. albicans* infection, and cytoplasmic HMGB1 became progressively elevated, maintaining a comparatively high level on days five and seven. Nuclear and cytoplasmic HMGB1 in the kidney was significantly increased after *C. albicans* infection. However, nuclear and cytoplasmic HMGB1 in the liver and kidney of the EP group were lower than the invasive *C. albicans* infection group, at the same time point. This observation indicates that HMGB1 also may transfer from the nucleus to the cytoplasm after *C. albicans* infection, with EP inhibiting this process. Similarly, EP also alleviated organ damage, improved mortality and inhibited TNF-α and IL-6 release. These results indicate that EP prevents lethality from invasive *C. albicans* infection by inhibiting HMGB1 expression and release.

In this study, we investigated the role of HMGB1 and the anti-HMGB1 effects of EP in invasive *C. albicans* infection. Our data suggests that HMGB1 plays an important role in invasive *C. albicans* infection and may provide a new effective target for the diagnosis and treatment of such infection. However, more clinical evidence is needed to precisely clarify this role. Additionally, further research is required to uncover HMGB1 molecular mechanisms in invasive *C. albicans* infection.

### Conclusions

Serum HMGB1 mRNA and protein levels in patients with severe sepsis with *C. albicans* infection are elevated and are positively correlated with PCT, CRP, BDG and disease severity. The combined detection of these indices may contribute to the early diagnosis of fungal sepsis. HMGB1 mRNA and protein levels were also significantly increased in mice with *C. albicans* infection. EP inhibited the expression and release of HMGB1, reducing tissue damage and mouse mortality, inhibiting TNF-α and IL-6 release, suggesting HMGB1 may play a vital role in invasive *C. albicans* infection and provide an effective therapeutic target for these infections.

## Acknowledgments

We acknowledge all the contributed authors. Chuanxin Wu and Jiaojiao Wang designed and performed the study, acquired and analyzed the data, and drafted the manuscript. Yunying Wang carried out the culture of *C. albicans*, and Chongxiang Chen performed the pathological analysis. Jing Chen and Xiaolong Rao selected the patients and collected specimens. Hang Sun designed and conceived this study and revised the manuscript critically. All authors read and approved the final manuscript.

## References

1. Wisplinghoff H, Bischoff T, Tallent SM, Seifert H, Wenzel RP, Edmond MB. 2014. Nosocomial bloodstream infections in US hospitals: analysis of 24,179 cases from a prospective nationwidesurveillance study. Clin Infect Dis 39:309–317.

2. Arendrup, Maiken C. 2010. Epidemiology of invasive candidiasis. Curr Opin Crit Care 16: 445–452.

3. Tragiannidis A, Tsoulas C, Kerl K, Groll AH. 2013. Invasive candidiasis: update on current pharmacotherapy options and future perspectives. Expert Opin Pharmacother 14:1515–1528.

4. Nieto MC, Tellería O, Cisterna R. 2015. Sentinel surveillance of invasive candidiasis in Spain: epidemiology and antifungal susceptibility. Diagn Microbiol Infect Dis 81:34–40.

5. De Pascale G, Tumbarello M. 2015. Fungal infections in the ICU: advances in treatment and diagnosis. Curr Opin Crit Care 21:421–429.

6. Méan M, Marchetti O, Calandra T. 2008. Bench-to-bedside review: Candida infections in the intensive care unit. Crit Care 12:204.

7. Drgona L, Khachatryan A, Stephens J, Charbonneau C, Kantecki M, Haider S, Barnes R. 2014. Clinical and economic burden of invasive fungal diseases in Europe: focus on pre-emptive and empirical treatment of Aspergillus and Candida species. Eur J Clin Microbiol Infect Dis 33:7–21.

8. Rodrigues ME, Silva S, Azeredo J, Henriques M. 2014. Novel strategies to fight Candida species infection. Crit Rev Microbiol 10:1–13.

9. Netea MG, Van Der Graaf CA, Vonk AG, Verschueren I, Van Der Meer JW, Kullberg BJ. 2002. The role of toll-like receptor(TLR)2 and TLR4 in the host defense against disseminated candidiasis. J Infect Dis 185:1483–1489.

10. Villamón E, Gozalbo D, Roig P, O’Connor JE, Fradelizi D, Gil ML. 2004. Toll-like receptor-2 is essential in murine defenses against Candida albicans infections. Microbes Infect 6:1–7.

11. Gil ML, Gozalbo D. 2006. TLR2, but not TLR4, triggers cytokine production by murine cells in response to Candida albicans yeasts and hyphae. Microbes Infect 8: 2299–2304.

12. Ferwerda G, Netea MG, Joosten LA, van der Meer JW, Romani L, Kullberg BJ. 2010. The role of Toll-like receptors and C-type lectins for vaccination against Candida albicans. Vaccine 28:614–622.

13. Akira S, Uematsu S, Takeuchi O. Pathogen recognition and innate immunity. 2006. Cell 124:783–801.

14. Beutler BA, Milsark IW, Cerami A. 1985. Cachectin/tumor necrosis factor: production, distribution, and metabolic fate in vivo. J Immunol 135:3972–3977.

15. Wang H, Bloom O, Zhang M, Vishnubhakat JM, Ombrellino M, Che J, Frazier A, Yang H. 1999. HMG-1 as a late mediator of endotoxin lethality in mice. Science 285:248–251.

16. Lin Y, Chen L, Li W, Fang J. 2015. Role of high-mobility group box-1 in myocardial ischemia/reperfusion injury and the effect of ethyl pyruvate. Exp Ther Med 9:1537–1541.

17. Gupta SK, Rastogi S, Prakash J, Joshi S, Gupta YK, Awor L, Verma SD. 2000. Anti-inflammatory activity of sodium pyruvate-a physiological antioxidant. Indian J Physiol Pharmacol 44:101–104.

18. Ullo L, Ochani M, Yang H, Tanovic M, Halperin D Yang R, Czura CJ, Fink MP, Tracey KJ. 2002. Ethyl pyruvate prevents lethalith in mice with established lethal sepsis and systemic inflammation. Proc Ncad Sci USA 99:12351–12356.

19. Dellinger RP, Levy MM, Rhodes A, Annane D, Gerlach H, Opal SM, Sevransky JE, Sprung CL, Douglas IS, Jaeschke R, Osborn TM, Nunnally ME, Townsend SR, Reinhart K, Kleinpell RM, Anqus DC, Deutschman CS, Machado FR, Rubenfeld GD, Webb S, Beale RJ, Vincent JL, Moreno R, Surviving Sepsis Campaign Guidelines Committee including The Pediatric Subgroup. 2012. Suriving Sepsis Campaign: international guidelines for management of severe sepsis and septic shock, 2012. Intensive Care Med 39:165–228.

20. Lionakis MS, Lim JK, Lee CC, Murphy PM. 2011. Organ-Specific Innate Immune Responses in a Mouse Model of Invasive Candidiasis. J Innate Immun 3:180–199.

21. Kang R, Chen R, Zhang Q, Hou W, Wu S, Cao L, Huang J, Yu Y, Fan XG, Sun X. 2014. HMGB1 in health and disease. Mol Aspects Med 40:1–116.

22. Li LC, Gao J, Li J. 2014. Emerging role of HMGB1 in fibrotic diseases. J Cell Mol Med 18:2331–9.

23. Wang H, Vishnubhakat JM, Bloom O, Bloom O, Zhang M, Ombrellino M, Sama A, Tracey KJ. 1999. Proinflammatory cytokines(tumor necrosis factor and interleukin 1) stimulate release of high mobility group protein-1 by pituicytes. Surgery 126:389–92.

24. Andersson U, Wang H, Palmblad K, Aveberger AC, Bloom O, Erlandsson-Harris H, Janson A. 2000. High mobility group 1 protein (HMG-1) stimulates proinflammatory cytokine synthesis in human monocytes. J Exp Med 192:565–70.

25. Wu C, Sun H, Wang H, Chi J, Liu Q, Guo H, Gong J. 2012. Evaluation of high mobility group box 1 protein as a presurgical diagnostic marker reflecting the severity of acute appendicitis. Scand J Trauma Resusc Emerg Med 20:61.

26. Wang XW, Karki A, Du DY, Zhao XJ, Xiang XY, Lu XQ. 2015. Plasma levels of high mogility group box 1 increase in patients with posttraumatic stress disorder after severe blunt chest trauma: a prospective cohort study. J Surg Res 193:308–15.

27. Singh A, Feng Y, Mahato N, Li J, Wu C, Gong J. 2015. Role of high-mobility group box 1 in patients with acute obstructive suppurative cholangitis-induced sepsis. J Inflamma Res 8:71–7.

28. Akira S, Uematsu S, Takeuchi O. 2006. Pathogen recognition and innate immunity. Cell 124:783–801.

29. Abramson N, Melton B. 2000. Leukocytosis: basics of clinical assessment. American family physician 62:2053–2066.

30. Pierrakos C, Vincent JL. 2010. Sepsis biomarkers: a review. Critical care 14:R15.

31. Müller B, Becker KL, Schächinger H, Rickenbacher PR, Huber PR, Zimmerli W, Ritz R. 2000. Calcitonin precursors are reliable markers of sepsis in a medical intensive care unit. Critical care medicine 28: 977–83.

32. Persat F, Ranque S, Derouin F, Michel-Nguyen A, Picot S, Sulahian A. 2008. Contribution of the (1-3)-beta-D-glucan assay for diagnosis of invasive fungal infections. J Clin Microbiol 46:1009–1013.

33. Jaijakul S, Vazquez JA, Swanson RN, Ostrosky-Zeichner L. 2012. (1,3)-β-D-glucan as a prognostic marker of treatment response in invasive candidiasis. Clin Infect Dis 55:521–6.

34. Karageorgopoulos DE, Vouloumanou EK, Ntziora F, Michalopoulos A, Rafailidis PI, Falagas ME. 2011. β-D-glucan assay for the diagnosis of invasive fungal infections: a meta-analysis. Clin Infect Dis 52:750–70.

35. Gil ML, Gozalbo D. 2006. TLR2, but not TLR4, triggers cytokine production by murine cells in response to Candida albicans yeasts and hyphae Microbes Infect. 8: 2299–2304.

36. Wu CX, Sun H, Liu Q, Guo H, Gong JP. 2012. LPS induces HMGB1 relocation and release by activating the NF-ΚB-CBP signal transdution pathway in the murine macrophage-like cell line RAW264.7. J Surg Res 175:88–100.

37. Wu CX, He LX, Guo H, Tian XX, Liu Q, Sun H. 2015. Inhibition effect of clycyrrhizin in lipopolysaccharide-induced high-mobility croup box 1 releasing and expression from RAW264.7 cells. Shock 43:412–21.

38. Magna M, Pisetstky DS. 2014. The role of HMGB1 in the pathogenesis of inflammatory and autoimmune disease. Mol Med 20:138–46.

